# Conditional Prediction of RNA Secondary Structure Using NMR Chemical Shifts

**DOI:** 10.1101/554931

**Authors:** Kexin Zhang, Aaron T. Frank

## Abstract

Inspired by methods that utilize chemical-mapping data to guide secondary structure prediction, we sought to develop a framework for using assigned chemical shift data to guide RNA secondary structure prediction. We first used machine learning to develop classifiers which predict the base-pairing status of individual residues in an RNA based on their assigned chemical shifts. Then, we used these base-pairing status predictions as restraints to guide RNA folding algorithms. Our results showed that we could recover the correct secondary folds for nearly all of the 108 RNAs in our dataset with remarkable accuracy. Finally, we assessed whether we could conditionally predict the structure of the model RNA, microRNA-20b (miR-20b), by folding it using folding restraints derived from chemical shifts associated with two distinct conformational states, one a free (apo) state and the other a protein-bound (holo) state. For this test, we found that by using folding restraints derived from chemical shifts, we could recover the two distinct structures of the miR-20b, confirming our ability to conditionally predict its secondary structure. A command-line tool for **C**hemical **S**hifts to **B**ase-**P**airing **S**tatus (CS2BPS) predictions in RNA has been incorporated into our CS2Structure Git repository and can be accessed via: https://github.com/atfrank/CS2Structure.

## INTRODUCTION

Like proteins, ribonucleic acids (or RNAs) play critical functional roles within cells, and like proteins, an RNA’s function is determined by its structure(1, 2, 3). RNAs, however, do not necessarily adopt a single structure. Instead, RNAs can adopt and interconvert between distinct conformational states that are kinetically linked to form a complex network of accessible states. Such networks enable RNAs to, for example, function as regulatory switches by responding to environmental stimuli such as changes in temperature or changes in ligand or metabolite concentration(4, 5, 6, 7, 8). Mapping the conformational landscapes of RNAs(4, 9), which consist of the set of conformational states accessible to an RNA, is crucial in unraveling the complex relationships between their sequence, their structure, and their function.

Determining the secondary structure of an RNA, under a specific set of physiochemical conditions, is a crucial first step in uncovering links between its sequence, its structure, and its function(10). *In silico* methods can be used to predict the secondary structure of an RNA from sequence by identifying the secondary structure that minimizes its folding free energy(11, 12, 13). However, the structure that an RNA adopts depends not only on its sequence but also on the physiochemical environment in which the RNA “resides.” Urgently needed are methods that can predict the RNA structure from sequence, *conditioned* on the physiochemical environment of an RNA.

Herein, we implemented and tested a framework for *conditionally* predicting the secondary structure of RNAs based on *assigned* chemical shift data(14, 15). Specifically, we trained a set of machine learning classifiers that take as input the *assigned* chemical shifts of individual residues in an RNA and then predict the base-pairing status of each residue. We then used these base-pairing predictions as restraints to guide the folding of the target RNA. We discovered that the secondary structures generated using our chemical shift derived base-pairing status predictions as restraints were more consistent with NMR-derived secondary structure models than the models generated without these restraints. Moreover, for the microRNA-20b (miR-20b), we were able to accurately predict its secondary structure *conditioned* on the available chemical shift data for two distinct conformational states.

Collectively, our results demonstrate that the information content in *assigned* chemical shift data can be leveraged to *conditionally* predict the secondary structure of an RNA by combining machine learning tools and existing structure prediction algorithms. With access to the *assigned* chemical shift fingerprints of individual conformational states of an RNA, the hybrid modeling approach described in this study could be used to generate a hypothetical map of its conformational landscape, which will be particularly powerful when some of these states correspond to difficult-to-characterize transient states(16, 17).

## MATERIALS AND METHODS

### Structure and Chemical Shift Dataset

For 115 RNAs, atomic NMR structures and NMR chemical shifts were downloaded from the Protein Data Bank (PDB: http://www.pdb.org) and the Biological Magnetic Resonance Data Bank (BMRB: http://www.bmrb.wisc.edu/), respectively. Next, LARMOR^D^(18) was used to predict chemical shifts using the coordinates of the first model in the NMR bundle for each RNA. Because ^13^C chemical shifts frequently contain systematic referencing errors(19), a structure-based approach was used to identify systematic referencing errors and, if necessary, correct ^13^C data for each RNA. Briefly, a Bayesian inference approach(20) was used to identify systematic offsets in the difference between experimental chemical shifts and chemical shifts computed using LARMOR^D^. For each type of non-exchangeable ^13^C nuclei (namely, C1’, C2’, C3’, C4’, C5’, C2, C5, C6, and C8), the corresponding chemical shift data were assumed to contain a systematic error if the mean estimated offset (*μ*_error_) was > 2 ppm and the ratio of the mean estimated offset and the standard deviation (*μ*_error_/*σ*_error_) was > 5. This approach was able to reproduce experimentally validated referencing errors previously identified by Aeschbacher *et* al.(19). The R code used to detect and correct the chemical shifts and the corrected dataset are available at: https://github.com/atfrank/CS2Structure.

After making any necessary corrections to the chemical shift data for each of the 115 RNAs in our initial dataset, we determined the weighted (or reduced) mean absolute error (wMAE) between the corrected experimental chemical shifts and chemical shifts computed from the first model of each of the NMR bundles. RNAs that exhibited ^1^H or ^13^C wMAE that were > 1.5 × the interquartile range (IQR), were considered outliers and removed from our dataset. The PDBIDs of the RNAs that were removed are: (1) 1S9S, (2) 1TJZ, (3) 2LC8, (4) 2AHT, (5) 2MQT, (6) 2M24, and (7) 5KMZ. For the remaining 108 RNAs, the secondary structures were retrieved using the program DSSR from the 3DNA suite(21, 22). The canonical base-pairing status of each residue in the RNA was determined from the secondary structure model.

### Chemical Shift Imputation

To impute missing chemical shift data, we used the R package MICE(23) with predictive mean matching (“pmm”). MICE assumes the probability of a data point being missing depends only on the observed data. It first fits a linear model for each variable (here, the chemical shifts of a specific nucleus type) based on all the other variables (the chemical shifts of the other nucleus types). Then, for a specific nucleus type, it identifies a set of observed chemical shifts whose imputed values are closest to the imputed value of the missing data point and randomly selects one of the observed values to fill in the missing data point. By building models on randomly selected subsets of data, multiple complete sets of chemical shifts are generated.

For each of the 108 RNA systems in our dataset, the imputed non-exchangeable (namely, C1’, C2’, C3’, C4’, C5’, C2, C5, C6, C8, H1’, H2’, H3’, H4’, H2, H5, H5’, H5”, H6, and H8) chemical shifts, along with the residue types and base-pairing status of individual residues were combined to produce a single data structure in which the rows corresponded to individual residues from each of the 108 RNAs and the columns corresponded to different nucleus types.

**Figure 1.**
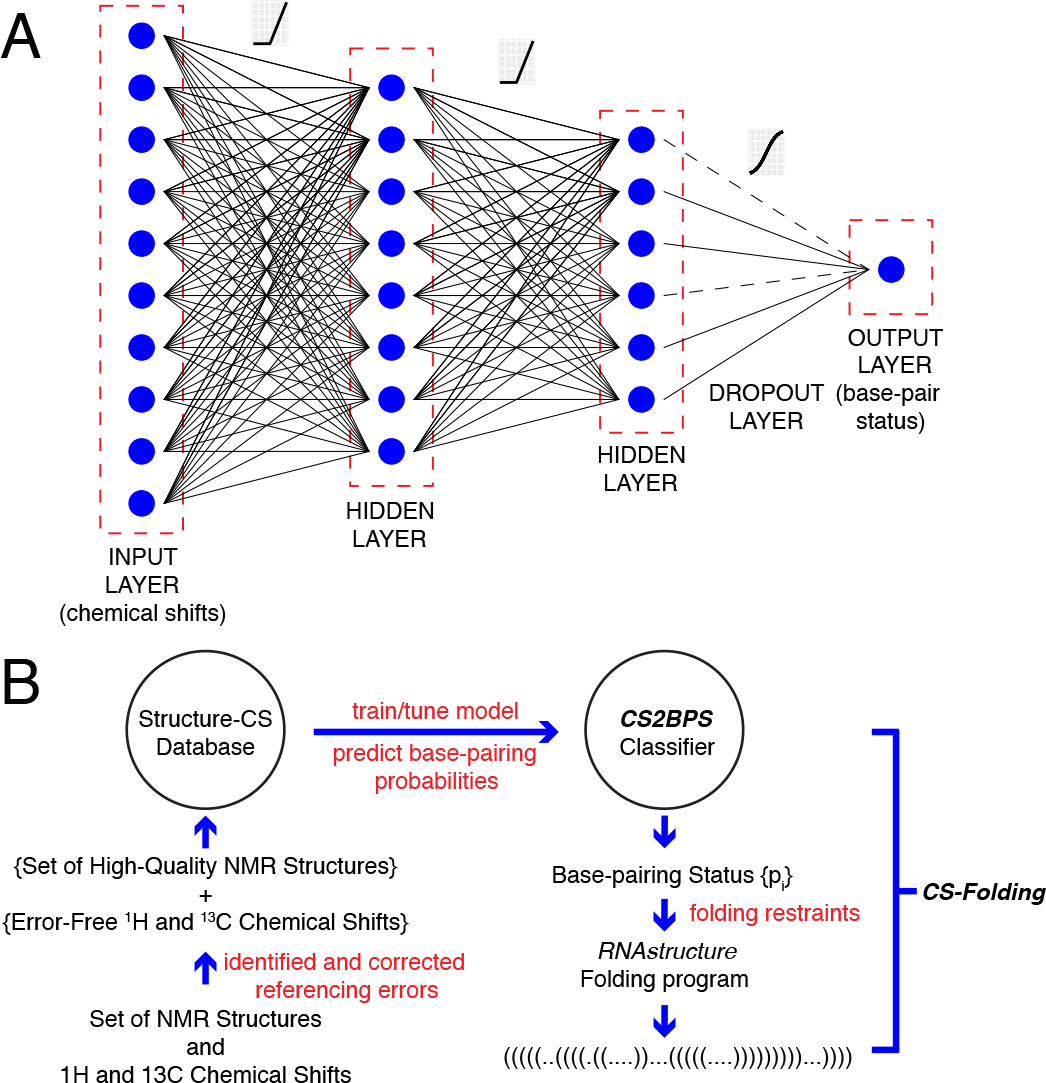
**A:** Artificial neural networks (ANNs). An ANN consist of an input layer, a set of hidden layers, and an output layer. We also added a dropout layer between the second hidden layer and the output layer, which randomly removed neural network connections (dashed lines), in order to avoid potential over-fitting. In the present work, we fed in chemical shifts associated with an RNA residue through the the input layer and the output layer, which had one neuron, returned the probability that the residue was not base-paired. For a given network architecture, the weights of connections between the neurons in each layer defined the ANN model. **B**: **C**hemical **S**hift to **B**ase-**B**airing **S**tatus (CS2BPS) workflow and CS-Folding framework. When developing CS2BPS classifiers, we first obtained a dataset containing NMR chemical shifts and NMR-derived secondary structures for 108 RNAs. Using a leave-one-out approach, independent sets of CS2BPS classifiers were trained and tuned for each RNA in our dataset via 5-fold cross validation. Next, in the CS-Folding framework, the optimized CS2BPS models were then applied on the target RNA to predict the base-pairing status of its individual residues. The average base-pairing status predictions calculated from independent sets of CS2BPS classifiers were then used as restraints in RNA folding simulations.

### Artificial Neural Network Classifiers

To predict the base-pairing status of individual residues in an RNA based on the observed chemical shifts of atoms in those residues, we trained a set of artificial neural network (ANN) classifiers(24). In an ANN, input features and output labels are connected through one or more layers of hidden neurons (Figure 1A). The neurons on adjacent layers are connected to each other through a linear transformation followed by an activation function. When training neural networks, gradient descent is performed to update network weights through back-propagation until a tolerable loss is achieved.

Here, we built a chemical shift-based ANN classifier (referred to hereafter as, **C**hemical **S**hift to **B**ase-**B**airing **S**tatus classifier, **CS2BPS**) consisting of two dense layers and one dropout layer. The dropout layer had the effect of regularizing the ANN and thus mitigating potential over-fitting. For a given residue *i*, the chemical shift data of residues *i*, *i*−1, and *i*+1, along with the residue types of *i*, *i−*1, and *i*+1 were fed into the CS2BPS classifier and the classifier outputted the probability of residue *i* being not base-paired (Figure 1B).

Hyper-parameters, namely, the number of neurons in each hidden layer, the loss function, the learning rate, the drop-out rate, the batch size, the optimization method and the number of epochs were optimized through grid search cross validation. The CS2BPS base-pairing status classifier was trained using Keras with a TensorFlow backend. We applied a ReLU activation function on both hidden layers and a sigmoid activaton function on the output layer (Figure 1A).

Using a leave-one-out approach, six independent CS2BPS classifiers were generated for each RNA in our dataset. For each RNA, RNAs in the training set with high sequence similarity (≥ 80%), determined using the tool vsearch(25), were first removed from the training set to avoid the “twinning” effect. Next, the set of six independent CS2BPS classifiers were generated and used to predict the base-pairing status of individual residues in the left-out (testing) RNA.

To quantify the accuracy of the classifiers, we computed the true positive rate (TPR), defined as the probability that a base-paired residue is predicted to be base-paired, and the true negative rate (TNR), defined as the probability that a not base-paired residue is predicted to be not base-paired, and the overall accuracy, defined as the fraction of residues in an RNA whose base-pairing status are correctly predicted, for each testing RNA. Here, the predicted base-pairing status corresponded to the average base-pairing probabilities calculated from those six CS2BPS classifiers (see above).

### Assessing the use of CS2BPS Classifiers to Guide Secondary Structure Prediction

To assess the use of the CS2BPS classifiers to guide secondary structure prediction, we implemented a CS-Folding framework. Within this CS-Folding framework, the base-pairing status predictions, which were calculated by averaging the results of six independent CS2BPS classifiers, were used as restraints in RNA folding simulations (Figure 1B). Similar to the approach used to incorporate SHAPE-derived restraints(26, 27, 28, 29) into RNA folding simulations, we utilized a restraint term of the form

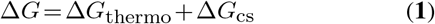

where Δ*G* is total folding free energy, Δ*G*_thermo_ is the thermodynamic free energy of folding, and Δ*G*_cs_ is the base-pairing restraint term which has the functional form

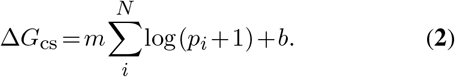

Here *m* and *b* are restraint parameters that influence the magnitude of the restraint free energy relative to the thermodynamic free energy, and *p*_*i*_ is the probability of a residue being not base-paired (Figure 1B) that is obtained from our CS2BPS classifiers.

We implemented the CS-Folding framework using folding algorithms from the RNAstructure modeling suite(30). Briefly, for each of the 108 RNAs in our dataset, CS-Folding simulations were carried out using the folding algorithms Fold(31), MaxExpect(32), and ProbKnot(33). Fold predicts secondary structures by free energy minimization. MaxExpect generates secondary structure models that contain highly probable base-pairs. And ProbKnot, like MaxExpect, generates secondary structure models that contain highly probable base-pairs, but allows pseudoknots. In all CS-Folding simulations, *m* and *b* were set to 1.8 kcal/mol and 0.6 kcal/mol, respectively. These values corresponded to the default values used to incorporate normalized SHAPE reactivities into RNA folding simulations(34). To select the best model among the Fold, MaxExpect, and ProbKnot generated structures with and without using CS2BPS-derived predictions as restraints, we defined the consistency score as the fraction of {*s*_fold_} that were identical to {*s*_CS2BPS_}, where {*s*_fold_} is the base-pairing status of individual residues from Fold, MaxExpect, and ProbKnot generated structures and {*s*_CS2BPS_} is the base-pairing status derived from the CS2BPS classifiers. Among these six possible structures, the one that has the highest consistency score with CS2BPS base-pairing predictions was selected as the predicted secondary structure model for a given RNA.

To compare the secondary structure models predicted using CS-Folding and the native secondary structures, we used the program scorer(11), from the RNAStructure suite. Given a reference secondary structure model and a comparison structure, scorer calculates the true positive rate (TPR), defined as the fraction of base-pairs in the comparison structure that also appear in the reference structure, and the positive predictive value (PPV), defined as the fraction of reference base-pairs that also appear in the comparison structure. In our case, the reference structure is the native NMR-derived structure and the comparison structure is the CS-Folding generated structure.

## RESULTS

### Model Assessment: Base-pairing status from chemical shifts

We began our study by training a set of machine learning classifiers to predict the base-pairing status of individual residues in an RNA from the ^1^H and ^13^C chemical shift “fingerprints” of each residue. To develop these classifiers, we utilized the artificial neural network machine learning technique (Figure 1A). Briefly, an artificial neural network (ANN) takes in information through an *input layer*, then passes it through one or more *hidden layers* and finally passes it through an *output layer*. Our ANN classifiers, which we refer to as **C**hemical **S**hift to **B**ase-**B**airing **S**tatus (**CS2BPS**) classifiers, are fed the chemical shifts for individual residues in an RNA as well as the chemical shifts of the residues before and after it. The network outputs the base-pairing status of each residues (i.e., the probability that a given residue is base-paired to some other residue). We note that for residues that are predicted to be base-paired, our CS2BPS classifiers *do not identify their base-pairing partner*.

Using a dataset containing NMR chemical shifts and NMR-derived secondary structures for 108 RNAs, we built six independent CS2BPS classifiers for each RNA in our dataset using a leave-one-out approach. Briefly, to build each classifier, data associated with one of the 108 RNAs (the left-out RNA) were removed from the dataset. Then six CS2BPS classifiers were trained using data from the other RNAs (i.e., the training set). The resulting CS2BPS classifiers were then tested on the left-out RNA. To mitigate bias due to the “twinning” effect in which data in training set closely resemble the left-out data, when building each CS2BPS classifier, we also excluded from the training set data associated with any RNA(s) that exhibited a high sequence similarity (≥ 80%) to the left-out RNA (see Methods).

Figure 2 summarizes the accuracy of the 108 CS2BPS classifiers. Reported are the true positive rate (TPR) and the true negative rate (TNR). Here a residue is base-paired if its mean classification probability was ≥ 0.4. This classification threshold was chosen so as to maximize the overall classification accuracy (that is, the fraction of residues whose base-pairing status were correctly predicted).

#### Overall Accuracy

The mean true positive rate (TPR) and the mean true negative rate (TNR) of the CS2BPS classifiers were 0.95 and 0.72, respectively; TPR values ranged between 0.77 and 1.00 (Figure 2A) and TNR values ranged between 0.00 and 1.00 (Figure 2B). These results indicate that our CS2BPS classifiers were better at identifying base-paired residues than unpaired residues. The comparatively lower TNR of our CS2BPS classifiers can be attributed to an imbalance in our dataset; the total number of base-paired and unpaired residues contained in our dataset were 2329 and 1023, respectively. This imbalance in our dataset might also explain why the classification threshold that maximized the overall classification accuracy was 0.4, rather than 0.5(35).

Out of the 108 systems in our dataset, 22 corresponded to systems for which only ^1^H chemical shifts were available. For these 22 systems with only ^1^H chemical shifts, and for which the corresponding ^13^C chemical shifts had to be imputed (see Methods), the mean TPR value was 0.93 (Figure 2A; *orange*); by comparison, the mean TPR value for systems for which both ^1^H and ^13^C chemical shifts were available was 0.96 (Figure 2A; *blue*). On the other hand, for systems for which only ^1^H chemical shifts were available, the mean TNR value was 0.67 (Figure 2B; *orange*), compared to 0.73 for systems for which both ^1^H and ^13^C chemical shifts were available (Figure 2B; *blue*). As might be expected, the CS2BPS classifiers exhibited higher TPR and TNR when both ^1^H and ^13^C chemical shifts were available.

#### Accuracy by residue types

Shown in Table 1 is a breakdown of the CS2BPS performance for individual residue types, namely G (guanine), C (cytosine), A (adenine), and U (uracil). The TPRs ranged between 0.90 and 0.97. For G and C residues, the TPRs were 0.96 and 0.97, respectively. By comparison, the TPRs for A and U residues were 0.90 and 0.92. The TNRs for individual residue types ranged between 0.56 and 0.80. For G and C residues, the TNRs were 0.66 and 0.56, respectively. By comparison, the TNRs for A and U residues were slightly higher: the values were 0.80 and 0.64, respectively.

#### Accuracy by base-pair types

Though our CS2BPS classifiers cannot predict the base-pairing partners for residues that are estimated to have a high probability of being base-paired, we were nonetheless interested in exploring whether our CS2BPS classifiers were able to correctly predict the base-pairing status of both residues in individual base-pairs. To explore this, all of the GC (or CG), AU (or UA), and GU (or UG) base-pairs were identified, and a TPR score was calculated for each base-pair type. In this case, the TPR was defined as the probability that both residues in a base-pair were correctly predicted to be base-paired. We found that for GC, AU, and GU base-pairs, the TPR values were 0.93, 0.88, and 0.75, respectively (Table 1). Examining the number of instances of each base-pair type in our database indicates these differences in TPR is most likely a result of an imbalance among the different types of base-pairs in our dataset: the total instances of GC base-pairs were 1374 compared to only 772 and 183 for AU and GU, respectively (Table 1).

**Table 1.** TPR and TNR by residue and base-pair type

| Residue Type | TPR | TNR | Instances |
| --- | --- | --- | --- |
| G | 0.96 | 0.66 | 985 |
| C | 0.97 | 0.56 | 859 |
| A | 0.90 | 0.80 | 749 |
| U | 0.92 | 0.64 | 759 |
| Base-pair type | TPR | TNR | Instances |
| GC | 0.93 | N/A | 1374 |
| AU | 0.88 | N/A | 772 |
| GU | 0.75 | N/A | 183 |

Here, TPR is defined as the probability that both residues in a base-pair are correctly predicted to be base-paired.

#### Representative examples

For the 47-nt Fluoride Riboswitch RNA (PDBID: 5KH8)(36), our CS2BPS predictions exhibited TPR and TNR values of 0.93 and 0.53, respectively (Figure 2C). This was one of the 4 structures in our dataset that contained pseudoknot interactions and whose overall prediction accuracy was only around the 9th percentile. Interestingly, for this RNA, most of the residues that participated in long-range tertiary contacts were correctly predicted to be base-paired (namely, residues 7-12 and 39-44) (Figure 2C). For this RNA, the major sources of error were due to the mis-classification of residues 35-38 and 45-47. Interestingly, for 5 out of the 8 mis-classified residues (namely, residue 4, 5, 35, 36, 37, 38, 42, and 44), we found significant variance in their classification across the six independent CS2BPS classifiers (Figure 2C; Table 4).

The second example, the 34-nt Simian Immunodeficiency Virus (SIV) RNA (PDBID: 2JTP)(37) was selected because its classification accuracy was identical to the the median classification accuracy (0.88) across the entire dataset. For this RNA, our CS2BPS predictions exhibited TPR and TNR values of 0.85 and 1.00, respectively (Figure 2D). The errors were due to residues 11, 15, 16, and 24 being mis-classified as not base-paired. As was the case for some of the mis-classified residues in the fluoride riboswitch, two of the mis-classified residues of the SIV RNA also exhibited high variance in their CS2BPS classification (Figure 2D; Table 5).

**Figure 2.**
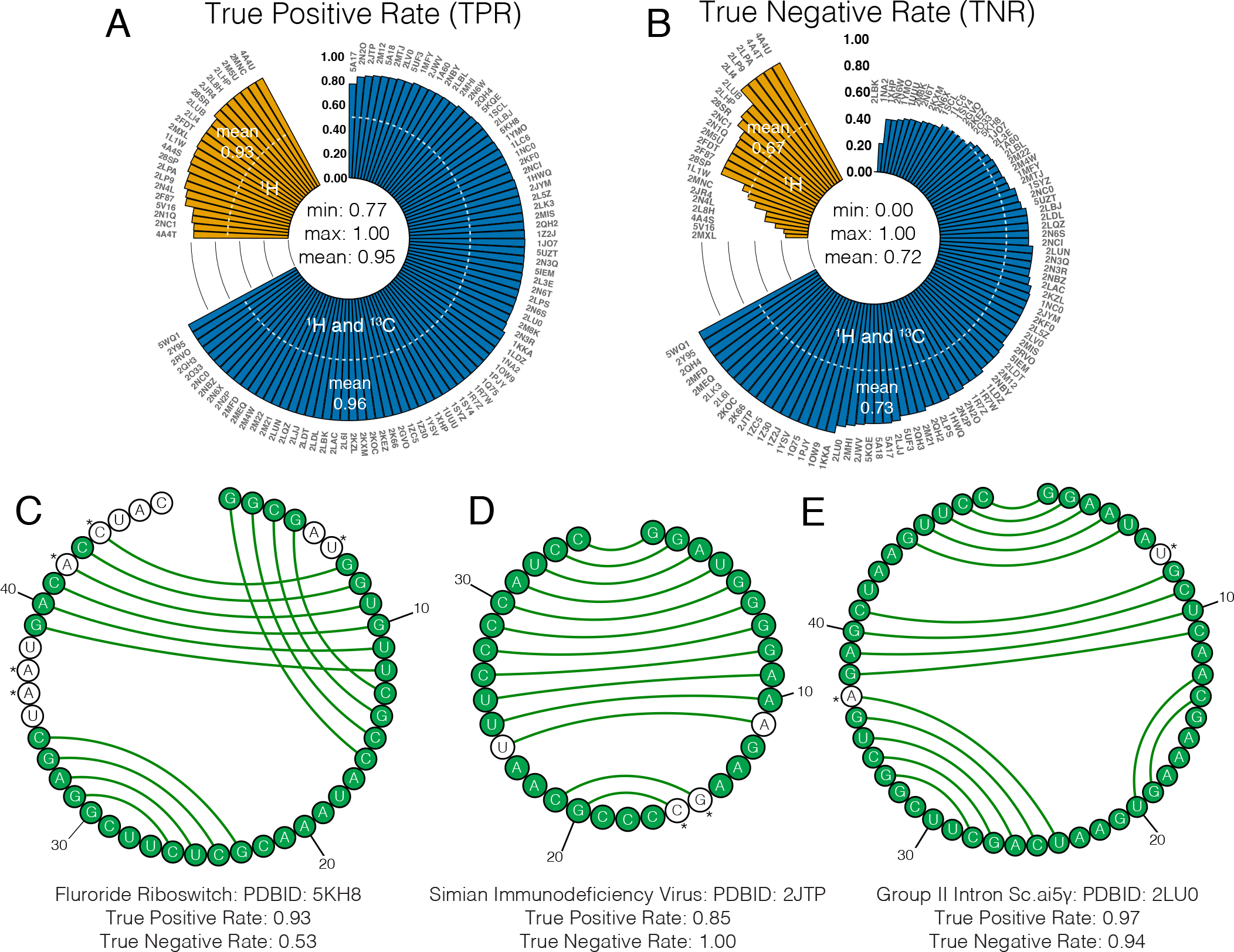
**Upper: CS2BPS Classification Accuracy.** Shown are circular barplots of the (A) true positive rate (TPR) (the fraction of residues that are correctly predicted as being base-paired) and (B) true negative rate (TNR) (the fraction of residues that are correctly predicted as being not base-paired). Accuracy statistics are based off a leave-one-out analysis in which one RNA system is removed from the training set, the remaining data used to train a classifier, and then the resulting classifier used to predict the base-pairing status of the left-out system. As a guide, the 0.5 accuracy levels are shown in white dashed lines. In the plots, bars are grouped based on whether only ^1^H (*orange*) or whether both ^1^H and ^13^C (*blue*) non-exchangeable chemical shifts were available in the corresponding RNA systems. **Lower: Visualizing CS2BPS results.** Shown are the CS2BPS predictions projected onto the native structures of (C) the Fluoride Riboswitch (PDBID: 5KH8), (D) the Simian Immunodeficiency Virus RNA (PDBID: 2JTP), and (E) the Group II Intron Sc.ai5*γ* RNA (PDBID: 2LU0). We also compared the base-pairing status prediction with the base-pairing status in the NMR structure: residues in green circles indicate that our CS2BPS predictions were consistent with the base-pairing status in the native structure while residues in white circles indicate that our CS2BPS predictions were incorrect. Residues labeled with ‘*’ exhibited high variance (see Table 4, 5, and 6) in their base-pairing classification across six independent CS2BPS classifiers.

For the third example, the 49-nt Group II Intro Sc.ai5*γ* RNA (PDBID: 2LU0)(38), our CS2BPS predictions exhibited TPR and TNR values of 0.97 and 0.94, respectively (Figure 2E). The only two residues that were mis-classified, residue 7 and 37, also exhibited high variance in their CS2BPS classification (Figure 2E; Table 6).

In general, we discovered that the TNR was significantly lower than the TPR for CS2BPS classifiers. In some cases, we found that a fraction of residues that were mis-classified exhibited high variance in their base-pairing predictions (See Table 4, 5, and 6). It should be noted that not all residues with high prediction variance were mis-classified. Collectively, however, these results show that, given a set of *assigned* ^1^H and ^1^3C chemical shifts for a given RNA, our CS2BPS classifiers could be used to predict the base-pairing status of individual residues.

### Guiding RNA secondary structure prediction

Next, we examined whether the residue-wise base-pairing probabilities that were predicted using our CS2BPS classifiers could be used to guide RNA secondary structure modeling. Given that an RNA can adopt distinct conformational states, each with a set of distinct chemical shift fingerprints, we sought to develop an approach that allowed us to predict the secondary structure of an RNA *conditioned* on a set of *assigned* chemical shift data. Such a method would be useful in mapping the structure landscape of an RNA from available chemical shift data.

#### CS-Folding: Chemical shift-derived base-pairing predictions as folding restraints

To predict RNA secondary structure *conditioned* on a set of chemical shifts, we implemented a CS-Folding framework in which CS2BPS-derived pairing predictions were used as restraints in RNA folding simulations (Figure 1B). Within this modeling framework, chemical shifts were taken as inputs and fed into a CS2BPS classifier to predict the base-pairing status of individual residues. These predictions were then used as restraints to guide RNA folding to produce a secondary structure model. This modeling approach closely resembles the approach used to guide modeling using chemical mapping data(28, 29).

Briefly, for each of the 108 RNAs in our dataset, we predicted secondary structure using the Fold, ProbKnot, and MaxExpect algorithms in RNAstructure suite, both with and without the single residue pairing restraints derived from the corresponding CS2BPS classifiers (Eq. **2**). Among these six predicted structures, the structure that was most consistent with the CS2BPS base-pairing predictions was selected and then compared to the NMR-derived reference secondary structure model.

**Table 2.** CS-Folding Accuracy

| Folding Algorithms | TPR | PPV |
| --- | --- | --- |
| Fold | 0.94/0.96 | 0.93/0.95 |
| ProbKnot | 0.95/0.96 | 0.92/0.93 |
| MaxExpect | 0.94/0.96 | 0.93/0.96 |
| **CS-Folding** | **0.97** | **0.95** |

TPR: is defined as the fraction of base-pairs in the predicted structure that also appeared in the NMR-derived structure; PPV: is defined as the fraction of base-pairs in the NMR-derived structure that also appeared in the predicted structure; TPR and PPV were calculated using the program scorer in the RNAstructure suite(11); For Fold, ProbKnot and MaxExpect, values obtained with and without using CS2BPS predictions as folding restraints are separated by '/'. Using Fold, ProbKnot and MaxExpect with and without CS2BPS predictions, we obtained six secondary structure models. Then the base-pairing status consistency scores between these six secondary structure models and CS2BPS predictions were calculated and the model with the highest consistency score was selected as the CS-Folding secondary structure model.

In Table 2, the results of the comparison between the final CS-Folding structures and the NMR-derived models are shown for the 108 systems in our dataset. Over the entire dataset, the mean TPR and PPV were 0.97 and 0.95, respectively. By comparison, when predicting secondary structures without the CS2BPS-derived base-pairing probabilities as restraints, the TPR and PPV were 0.94 and 0.93, 0.95 and 0.92, and 0.94 and 0.93, using Fold, ProbKnot, and MaxExpect, respectively.

Shown in Figure 3 are circular barplots of the TPRs and PPVs for the CS-Folding results of the 108 RNAs in our dataset. In general, the CS-Folding generated structures exhibited high TPR (0.97) and high PPV (0.95) values. For RNAs for which only ^1^H chemical shifts were available, the prediction TPR and PPV were 0.98 and 0.99, respectively (Figure 3A and 3B). In comparison, for RNAs for which both ^1^H and ^1^3C chemical shifts were available, the TPR and PPV values were 0.96 and 0.94, respectively (Figure 3A and 3B).

**Table 3.** CS-Folding TPR by base-pair type

| Base-pair Type | TPR | Instances |
| --- | --- | --- |
| GC | 0.97 | 1378* |
| AU | 0.98 | 772 |
| GU | 0.90 | 184* |

Shown in Table 3 are the TPRs for the recovery of GC, AU and GU base-pairs in the CS-Folding generated structures. In general, CS-Folding framework was able to recover all three base-pair types with high TPRs, 0.97, 0.98 and 0.90 for GC, AU and GU base-pairs, respectively. The slightly lower TPR for GU base-pairs was most likely due to the lower instances of GU base-pairs in the 108 RNAs dataset. Our dataset contained 184 of GU base-pairs compared to 772 AU and 1378 GC base-pairs, respectively.

Shown in Figure 3C, 3D, and 3E are detailed comparisons between the native secondary structures and the CS-Folding results for the Fluoride Riboswitch RNA (PDBID: 5KH8) (Figure 3C), the SIV RNA (PDBID: 2JTP) (Figure 3D), and the Group II Intron Sc.ai5*γ* RNA (PDBID: 2LU0)(Figure 3E), respectively. For each structure, only the canonical bases pairs are shown.

For the Fluoride Riboswitch RNA, the TPR was 0.93 (Figure 3C). The predicted CS-Folding structure recovered 5 out of the 6 pseudoknotted base-pairs as well as all of the non-pseudoknotted base-pairs. Interestingly, for the pseudoknotted U12-G39 base-pair that was missing in the CS-Folding structure, our CS2BPS classifier predicted both of these residues to be base-paired (Figure 3C). For this RNA, the PPV was also high (0.93); the CS-Folding structure only contained a single extraneous A5-U12 base-pair (Figure 3C).

For the SIV RNA and the Group II Intron Sc.ai5*γ* RNA, the majority of the base-pairs in the reference NMR models were correctly recovered, 11 out of 13 (TPR=0.85)(Figure 3D) and 16 out of 16 (TPR= 1.00)(Figure 3E), respectively. In both cases, no extraneous base-pairs were found in the CS-Folding structures (PPV=1.00).

#### Application of CS-Folding to the microRNA-20b pre-element

We concluded our study by applying the CS-Folding framework to the microRNA-20b (miR-20b) pre-element, for which two distinct conformational states have recently been characterized using NMR spectroscopy: an unbound (*apo*) state (Figure 4A) and a Rbfox RRM protein-bound (*holo*) state (Figure 4B)(39). When interacting with the conserved Rbfox RRM protein, two canonical base-pairs that are present in the *apo* state are disrupted (Figure 4A and B), enabling the protein and the RNA to interact in a sequence-specific manner. In addition to atomic structures, the *assigned* chemical shift data corresponding to the *apo* and *holo* conformational states are available, enabling us to test whether we could use the two sets of *assigned* chemical shifts to *conditionally* predict structures of the miR-20b RNA. Using the *apo* and *holo* chemical shifts, we predicted the base-pairing status of each residue in miR-20b using our CS2BPS classifiers and then used the CS-Folding framework to predict their secondary structures.

**Figure 3.**
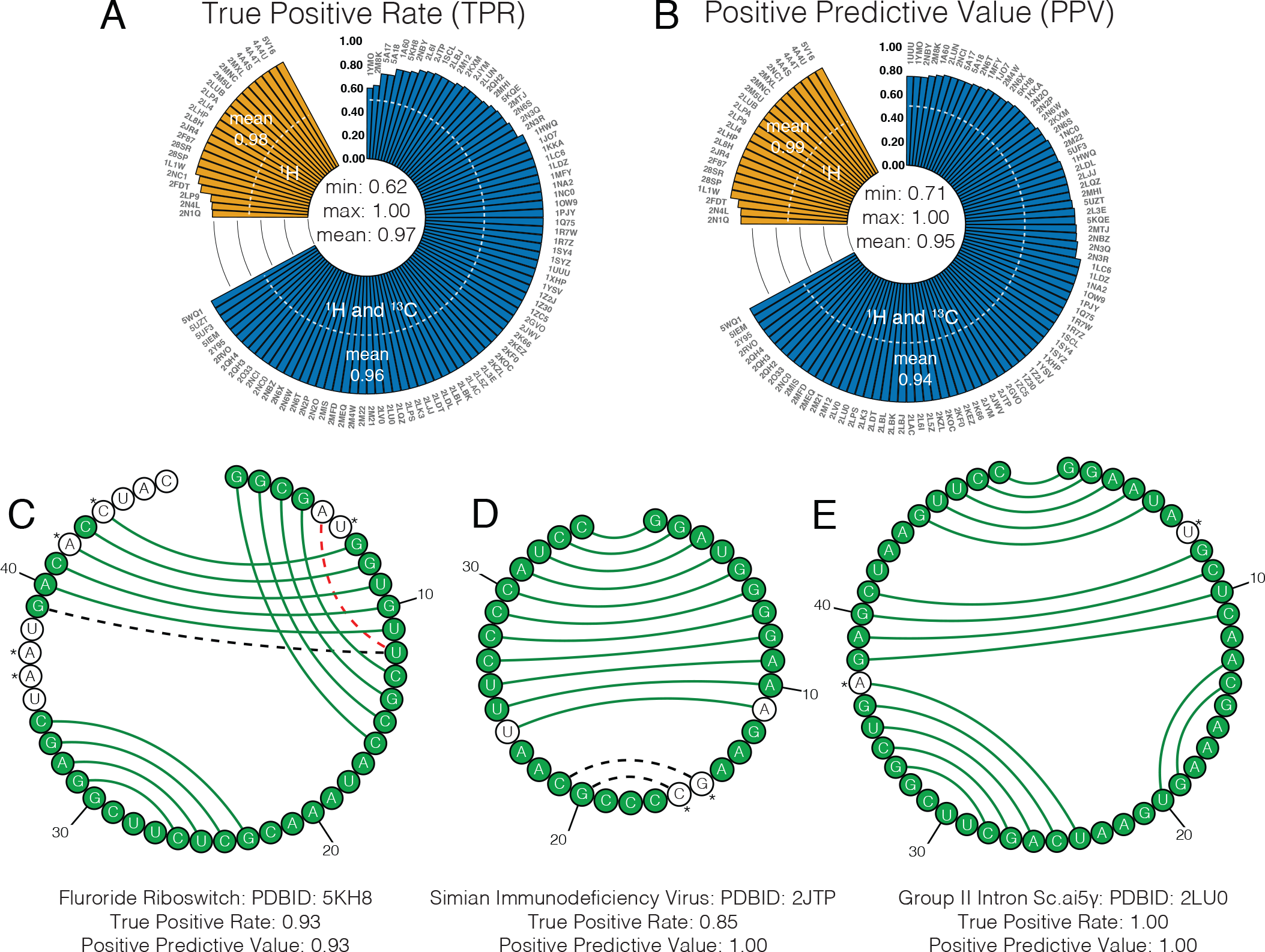
**Upper: CS-Folding Accuracy.** Shown are circular barplots of the (A) true positive rate (TPR) (the fraction of base-pairs in the CS-Folding structure that also appear in the NMR-derived structure) and (B) positive predicted value (PPV) (the fraction of base-pairs in the NMR-derived structure that also appear in the CS-Folding generated structure) obtained when comparing the reference NMR secondary structure of each RNA to the model obtained from folding the RNA using CS2BPS-derived base-pairing probabilities as folding restraints. For each RNA, the CS2BPS derived base-pairing probabilities were used as folding restraints to generate a model using the program Fold, ProbKnot and MaxExpect, respectively. Then the consistencies of the base-pairing status between the folded structures and the CS2BPS-derived predictions were calculated and the model with the highest consistency was selected. In the case where more than one structure had same consistency, the reference folding free energy of the secondary structure model was used to select the best model. As a guide, the 0.5 accuracy levels are shown in white dashed lines. In the plots, bars are grouped based on whether only ^1^H (*orange*) or whether both ^1^H and ^13^C (*blue*) non-exchangeable chemical shifts were available in the corresponding RNA systems. **Lower: Visualizing CS-Folding results.** Shown are the comparisons between CS-Folding predicted structures and secondary structure models derived from NMR bundle for (C) the Fluoride Riboswitch (PDBID: 5KH8), (D) the Simian Immunodeficiency Virus RNA (PDBID: 2JTP), and (E) the Group II Intron Sc.ai5*γ* RNA (PDBID: 2LU0). We also compared the base-pairing status prediction with the base-pairing status in the NMR structure: residues in green circles indicate that our CS2BPS predictions were consistent with the base-pairing status in the native structure while residues in white circles indicate that our CS2BPS predictions were incorrect. Base-pairs connected by green lines were present in both CS-Folding structure and NMR structure, while base-pairs connected by red dashed lines were only present in CS-Folding structure and base-pairs connected by black dashed lines were only present in NMR structure. Residues labeled with ‘*’ exhibited high variance (see Table 4, 5, and 6) in their base-pairing classification across six independent CS2BPS classifiers.

Shown in Figure 4A and 4B are the detailed comparisons between the native secondary structures and the predicted structures. In the case of the *apo* state, the TPR and PPV between native and predicted CS-Folding structures were both 1.00 (Figure 4A), indicating that we were able recover the native secondary structure. Similarly, for the *holo* state, the TPR and PPV between native and predicted CS-Folding structures were 1.00 and 0.80, respectively (Figure 4B). The only error in the CS-Folding generated structure of the *holo* state was an extra base-pair between residues G1 and C23. These results indicate that by biasing the folding algorithms using CS2BPS predictions, we were able to *conditionally* predict the two distinct conformational states of the miR-20b RNA.

**Figure 4.**
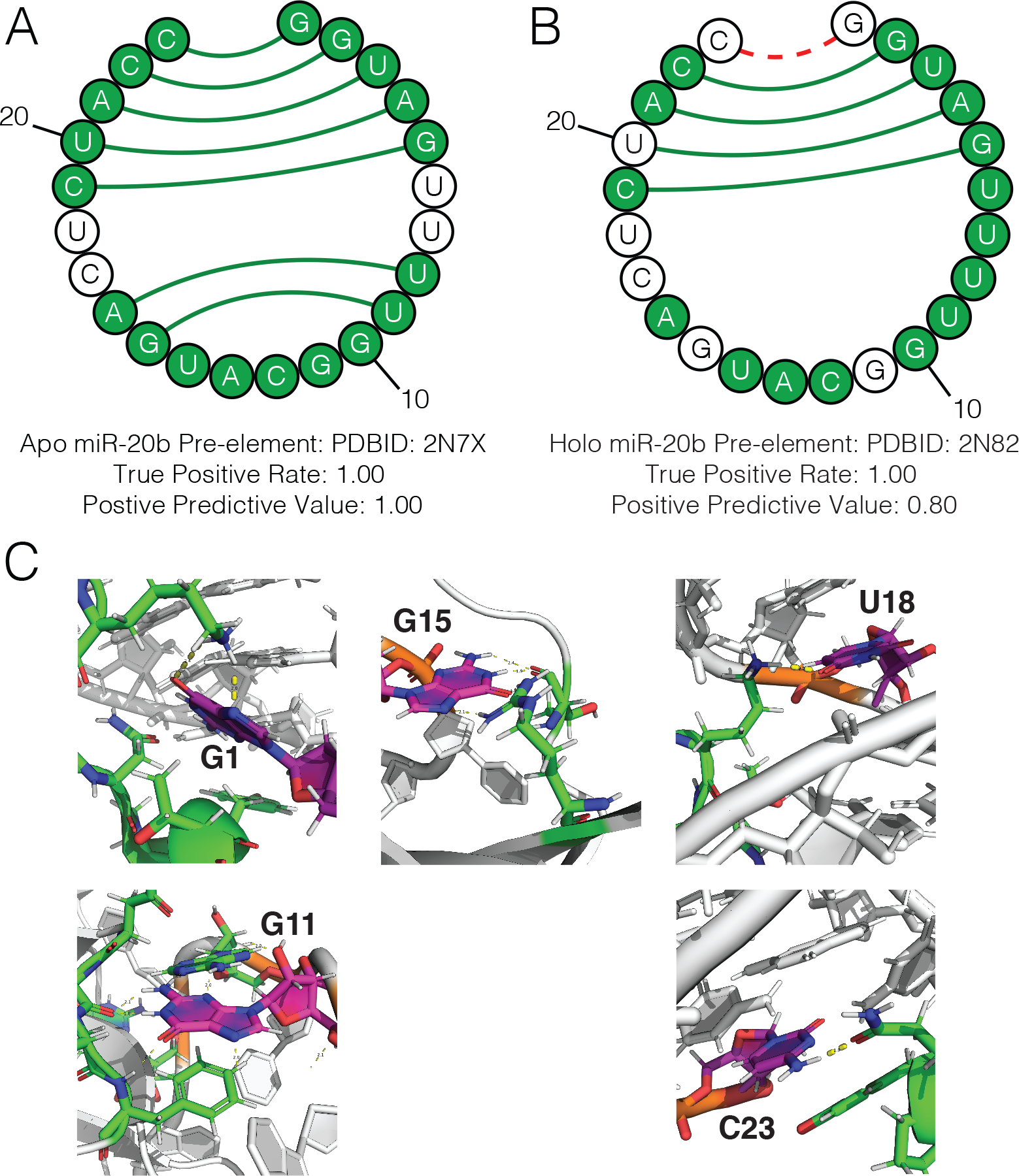
Visualizing CS-Folding results for the microRNA-20b (miR-20b) pre-element. Shown in **A** and **B** are the CS-Folding results of the *apo* and *holo* states of miR-20b, respectively. Shown in **C** are residues G1, G11, G15, G18, and U23, which, based on the secondary structure of the *holo* state of miR-20b were initially thought to be “mis-classfied” as being base-paired. However, closer examination of the 3D structure of the miR-20b-Rbfox complex revealed that these residues formed hydrogen bonds to residues in Rbfox.

Though the CS-Folding structures closely resembled the reference NMR structures of *apo* and *holo* states, respectively, the CS2BPS predictions which we used as folding restraints to predict their structures, contained what appeared, initially, to be several inconsistencies. For example, in the *apo* state, residues U6, U7, C16, and U17 were “mis-classified” as being base-paired (Figure 4A). Closer examination of the structure of the *apo* state (PDBID: 2N7X) revealed that these residues were, however, involved in non-canonical base-pairs, which we ignored in the study, because of their under-representation in our dataset and due to the fact that Fold, MaxExpect, and ProbKnot algorithms currently only predict canonical base pairs. Similarly, in the *holo* state, residues G1, G11, G15, C17, U18, U20 and C23 were all “mis-classified” as being base-paired, on the basis of the *holo* state secondary structure of the miR-20b RNA (Figure 4B). Closer examination of the structure of the *holo* state (PDBID: 2N82) (including the Rbfox RRM protein) revealed that with the exception of C17 and U20, these residues were involved in base-pairing interactions with the Rbfox RRM protein (Figure 4C).

## DISCUSSION

In this study, we generated a set of artificial neural network classifiers that were capable of predicting the base-pairing status of individual residues in RNAs directly from their non-exchangeable ^1^H and ^13^C chemical shift signature. These classifiers, which we referred to as CS2BPS (**C**hemical **S**hift to **B**ase-**P**airing **S**tatus) classifiers, were able to identify base-paired residues with high true positive rate (TPR), regardless of the residue type (Table 1). Interestingly, we found that even if only ^1^H chemical shifts were available, the base-pairing status of residues could still be accurately predicted (Figure 2A and 2B).

When assessing the accuracy of our CS2BPS-derived predictions for individual residues in base-pairs (Table 1), we discovered that, compared to the residues in GC and AU base-pairs, our CS2BPS classifiers exhibited poorer performance for individual Gs and Us in GU base-pairs (the TPR value was 0.75). This reduced accuracy was most likely due to the relatively small number of GU base-pairs (183) in our dataset, compared to GC base-pairs (1374) and AU base-pairs (772) (Table 1).

Heavily inspired by previous work in which single nucleotide SHAPE reactivities were used to guide RNA folding algorithms(28, 29), we explored whether the base-pairing status predictions derived from our CS2BPS classifiers could be used to guide RNA secondary structure folding. Within what we refer to as a CS-Folding framework, in which the CS2BPS-derived base-pairing status predictions (represented as single-residue pairing probabilities) were used as folding restraints (Eq. **1** and **2**), we found that we could recover the correct fold of most of the 108 RNAs in our dataset with remarkable accuracy. When guiding algorithms Fold, Probknot, and MaxExpect from RNAstructure suite with our CS2BPS-derived predictions and then identifying the structure with the highest base-pairing status consistency with our CS2BPS predictions, we were able to achieve mean TPR and PPV values of 0.97 and 0.95, respectively (Figure 3 and Table 2). By comparison, the TPR and PPV values we obtained when using Fold, ProbKnot and MaxExpect by themselves (that is, not restrained using our CS2BPS predictions) were 0.94 and 0.93, 0.95 and 0.92, 0.94 and 0.93, respectively (Table 2). Within the CS-Folding modeling framework, we discovered that our recovery of GU base-pairs was slightly lower than our recovery of GC and AU base-pairs; the mean TPR value for GU base-pairs was 0.90 compared to values of and 0.98 for GC and AU base-pairs respectively (Table 3). The slightly lower recovery rate for GU was, again, most likely due to the fact that the CS2BPS predictions for Gs and Us in GU base-pairs were less reliable (Table 1).

To test whether we could *conditionally* predict the secondary structure of an RNA, we applied our CS-Folding approach to microRNA-20b (miR-20b). For this RNA, two distinct conformational states, the free (*apo*) state and protein bound (*holo*) state, were recently characterized using NMR spectroscopy(39). Access to the structures and chemical shift data associated with both states of miR-20b enabled us to test whether we could use the CS-Folding framework to *conditionally* predict its secondary structure. We discovered that we could recover, with high TPR and PPV, the canonical base-pairs for the *apo* and *holo* conformational states of miR-20b, respectively (Figure 4A and 4B). This result suggests that given the chemical shifts for individual conformational states of an RNA, the CS-Folding modeling approach might be a viable technique for predicting the structure of each conformational state.

One significant limitation of our method is that it requires *assigned* chemical shifts. Indeed, when chemical shifts have been *assigned* for an RNA, base-pairing interactions within the RNA (and thus the secondary structure) can be directly determined using NOESY NMR experiments(40). The CS-Folding does not, therefore, provide a significant advantage over conventional NMR methods for determining the secondary structure of RNAs. However, we envision that the CS-Folding framework we demonstrated in this work can be used as a tool to *independently* validate NOSEY-derived secondary structural models of RNAs. More intriguing, though, is the possibility of using the CS-Folding framework to model the secondary structure of transient states of RNAs. Increasingly, there is keen interest in characterizing the transient states of RNAs. Unfortunately, it is not currently possible to detect the NOEs associated with these transient states. As such, conventional methods cannot be used to infer the secondary structure associated with the transient state or states of an RNA. Fortunately, it is now possible to characterize the ^1^H and ^13^C chemical shift signature of RNA transient states using techniques based on saturation transfer(36, 41) and relaxation dispersion(16, 42). The results we presented for the miR-20b RNA, suggest that with access to these ^1^H and ^13^C chemical shifts, a CS-Folding framework, which utilizes predictions derived from CS2BPS classifiers like the ones we developed in this work, could be used to generate putative models for the transient states of RNAs.

To facilitate the community-wide use of our CS2BPS classifiers, we have made a command-line tool available to the academic community with which users can predict the base-pairing status of individual residues in an RNA from their *assigned* chemical shifts. These CS2BPS predictions can then be used to guide RNA secondary structure prediction using external tools like Fold, ProbKnot, and MaxExpect (from the RNAStructure suite) or other RNA folding tools that accept and incorporate single residue pairing probabilities as folding restraints. The command-line tool has been incorporated into our CS2Structure repository and can be accessed via: https://github.com/atfrank/CS2Structure.

**Table 4.**
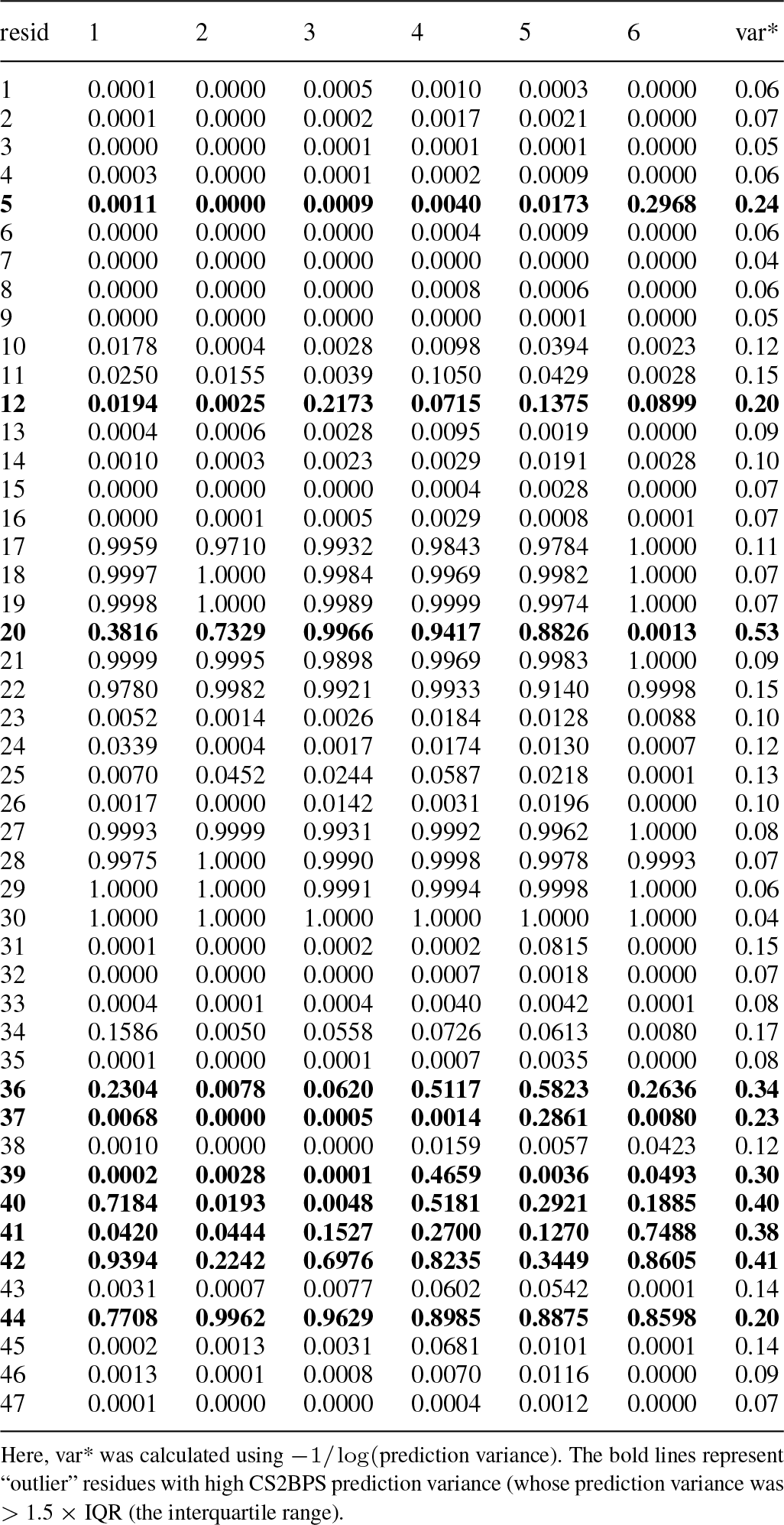
CS2BPS predictions for the Fluoride Riboswitch (PDBID: 5KH8)

**Table 5.** CS2BPS predictions for the Simian Immunodeficiency Virus (SIV) RNA (PDBID: 2JTP)

| resid | 1 | 2 | 3 | 4 | 5 | 6 | var* |
| --- | --- | --- | --- | --- | --- | --- | --- |
| 1 | 0.0015 | 0.0010 | 0.0056 | 0.0096 | 0.0055 | 0.0001 | 0.09 |
| 2 | 0.0553 | 0.0076 | 0.0114 | 0.0242 | 0.0092 | 0.0070 | 0.13 |
| 3 | 0.0721 | 0.0449 | 0.0891 | 0.0976 | 0.0678 | 0.1071 | 0.13 |
| 4 | 0.0018 | 0.0006 | 0.0106 | 0.0180 | 0.0092 | 0.0028 | 0.10 |
| 5 | 0.0021 | 0.0006 | 0.0036 | 0.0100 | 0.0071 | 0.0013 | 0.09 |
| 6 | 0.0008 | 0.0000 | 0.0007 | 0.0013 | 0.0006 | 0.0005 | 0.06 |
| 7 | 0.0016 | 0.0000 | 0.0032 | 0.0116 | 0.0015 | 0.0026 | 0.09 |
| 8 | 0.0217 | 0.0023 | 0.0178 | 0.0346 | 0.0088 | 0.0050 | 0.11 |
| 9 | 0.0048 | 0.0002 | 0.0272 | 0.0112 | 0.0048 | 0.0025 | 0.11 |
| 10 | 0.0043 | 0.0016 | 0.0220 | 0.0598 | 0.0071 | 0.0037 | 0.13 |
| 11 | 0.9373 | 0.9870 | 0.8510 | 0.8956 | 0.9090 | 0.9304 | 0.16 |
| **12** | **0.9630** | **0.9909** | **0.7291** | **0.8032** | **0.8493** | **0.4831** | **0.30** |
| **13** | **0.9608** | **0.8440** | **0.4732** | **0.5568** | **0.7560** | **0.4809** | **0.32** |
| 14 | 0.8537 | 0.9873 | 0.6945 | 0.7748 | 0.8013 | 0.8840 | 0.22 |
| **15** | **0.7419** | **0.5396** | **0.7312** | **0.4204** | **0.5434** | **0.8502** | **0.27** |
| **16** | **0.4963** | **0.9805** | **0.5220** | **0.4225** | **0.3977** | **0.9040** | **0.37** |
| 17 | 0.9962 | 0.9997 | 0.9962 | 0.9975 | 0.9982 | 0.9957 | 0.08 |
| 18 | 0.9861 | 0.9996 | 0.9931 | 0.9936 | 0.9942 | 0.9971 | 0.09 |
| 19 | 0.9718 | 0.9979 | 0.9923 | 0.9990 | 0.9969 | 0.9774 | 0.11 |
| 20 | 0.0157 | 0.0792 | 0.0220 | 0.0087 | 0.0118 | 0.0213 | 0.14 |
| 21 | 0.0105 | 0.0033 | 0.0227 | 0.3151 | 0.1347 | 0.0113 | 0.24 |
| 22 | 0.9912 | 0.9998 | 0.9893 | 0.9854 | 0.9913 | 0.8049 | 0.19 |
| 23 | 0.7439 | 0.8314 | 0.8475 | 0.7549 | 0.6228 | 0.7294 | 0.20 |
| 24 | 0.8946 | 0.9819 | 0.8687 | 0.9284 | 0.9243 | 0.9144 | 0.15 |
| **25** | **0.2970** | **0.4055** | **0.2289** | **0.1883** | **0.1647** | **0.5186** | **0.25** |
| 26 | 0.0269 | 0.0059 | 0.1526 | 0.0655 | 0.0597 | 0.0393 | 0.17 |
| 27 | 0.0144 | 0.0036 | 0.0229 | 0.0173 | 0.0142 | 0.0674 | 0.13 |
| 28 | 0.0106 | 0.0011 | 0.0372 | 0.0148 | 0.0146 | 0.0271 | 0.11 |
| 29 | 0.0181 | 0.0042 | 0.0678 | 0.0181 | 0.0197 | 0.0164 | 0.13 |
| 30 | 0.1026 | 0.0207 | 0.1014 | 0.0956 | 0.0718 | 0.0223 | 0.15 |
| 31 | 0.1394 | 0.0071 | 0.1098 | 0.0323 | 0.2332 | 0.0284 | 0.20 |
| 32 | 0.0065 | 0.0004 | 0.0208 | 0.0181 | 0.0115 | 0.0061 | 0.10 |
| 33 | 0.0014 | 0.0002 | 0.0091 | 0.0093 | 0.0023 | 0.0027 | 0.09 |
| 34 | 0.0263 | 0.0009 | 0.0323 | 0.0225 | 0.0334 | 0.0124 | 0.11 |

Here, var\* was calculated using $-1/\log(\text{prediction variance})$ . The bold lines represent “outlier” residues with high CS2BPS prediction variance (whose prediction variance was $> 1.5 \times \text{IQR}$ (the interquartile range)).

**Table 6.** CS2BPS predictions for the Group II Intron Sc.ai5*γ* RNA (PDBID: 2LU0)

| resid | 1 | 2 | 3 | 4 | 5 | 6 | var* |
| --- | --- | --- | --- | --- | --- | --- | --- |
| 1 | 0.0019 | 0.0071 | 0.0021 | 0.0006 | 0.0000 | 0.0021 | 0.08 |
| 2 | 0.0000 | 0.0000 | 0.0003 | 0.0003 | 0.0000 | 0.0003 | 0.06 |
| 3 | 0.0000 | 0.0000 | 0.0006 | 0.0024 | 0.0000 | 0.0004 | 0.07 |
| 4 | 0.1217 | 0.0074 | 0.1288 | 0.2312 | 0.0259 | 0.2124 | 0.21 |
| **5** | **0.3120** | **0.7393** | **0.4139** | **0.4779** | **0.1860** | **0.4989** | **0.30** |
| **6** | **0.9402** | **0.7611** | **0.5340** | **0.8485** | **0.9698** | **0.9211** | **0.28** |
| **7** | **0.2835** | **0.4900** | **0.5154** | **0.9249** | **0.6068** | **0.6851** | **0.33** |
| **8** | **0.1511** | **0.0571** | **0.1356** | **0.4864** | **0.0001** | **0.1577** | **0.28** |
| 9 | 0.0003 | 0.0001 | 0.0142 | 0.0031 | 0.0000 | 0.0047 | 0.10 |
| **10** | **0.1476** | **0.9891** | **0.4186** | **0.2001** | **0.0000** | **0.2334** | **0.48** |
| 11 | 0.0307 | 0.0015 | 0.1826 | 0.1690 | 0.0050 | 0.1927 | 0.21 |
| 12 | 0.9992 | 0.9995 | 0.9797 | 0.9459 | 1.0000 | 0.9756 | 0.13 |
| 13 | 0.0330 | 0.0262 | 0.1051 | 0.0246 | 0.1489 | 0.2349 | 0.20 |
| **14** | **0.4399** | **0.4969** | **0.1999** | **0.0375** | **0.0001** | **0.2801** | **0.31** |
| 15 | 1.0000 | 1.0000 | 0.9998 | 0.9973 | 1.0000 | 0.9955 | 0.08 |
| 16 | 1.0000 | 1.0000 | 1.0000 | 0.9986 | 1.0000 | 0.9972 | 0.07 |
| 17 | 0.9999 | 0.9999 | 0.9915 | 0.9877 | 1.0000 | 0.9862 | 0.10 |
| 18 | 1.0000 | 1.0000 | 1.0000 | 0.9994 | 1.0000 | 0.9997 | 0.06 |
| 19 | 0.0002 | 0.0001 | 0.0085 | 0.0004 | 0.0000 | 0.0036 | 0.09 |
| 20 | 0.0761 | 0.0510 | 0.3046 | 0.1181 | 0.0000 | 0.1899 | 0.23 |
| 21 | 0.9438 | 0.9967 | 0.8031 | 0.9051 | 1.0000 | 0.7763 | 0.21 |
| 22 | 0.9759 | 0.9935 | 0.9817 | 0.9686 | 0.9999 | 0.9475 | 0.13 |
| 23 | 0.9926 | 0.9513 | 0.8415 | 0.9305 | 0.9792 | 0.8993 | 0.17 |
| 24 | 0.0019 | 0.1350 | 0.0813 | 0.0312 | 0.1470 | 0.3329 | 0.23 |
| 25 | 0.0033 | 0.0006 | 0.0180 | 0.0104 | 0.0000 | 0.0274 | 0.11 |
| 26 | 0.0014 | 0.0005 | 0.0020 | 0.0022 | 0.0000 | 0.0033 | 0.07 |
| 27 | 0.0001 | 0.0002 | 0.0028 | 0.0004 | 0.0000 | 0.0015 | 0.07 |
| 28 | 0.0001 | 0.0000 | 0.0052 | 0.0027 | 0.0000 | 0.0055 | 0.08 |
| 29 | 0.9064 | 0.9965 | 0.9598 | 0.9475 | 1.0000 | 0.8572 | 0.17 |
| 30 | 0.9988 | 1.0000 | 0.9974 | 0.9871 | 1.0000 | 0.9882 | 0.10 |
| 31 | 1.0000 | 1.0000 | 0.9999 | 0.9947 | 1.0000 | 0.9991 | 0.08 |
| 32 | 1.0000 | 1.0000 | 1.0000 | 0.9882 | 1.0000 | 0.9985 | 0.09 |
| 33 | 0.0000 | 0.0000 | 0.0002 | 0.0002 | 0.0000 | 0.0000 | 0.05 |
| 34 | 0.0000 | 0.0000 | 0.0005 | 0.0003 | 0.0000 | 0.0025 | 0.07 |
| 35 | 0.0000 | 0.0001 | 0.0011 | 0.0026 | 0.0000 | 0.0020 | 0.07 |
| 36 | 0.0000 | 0.0000 | 0.0009 | 0.0043 | 0.0000 | 0.0007 | 0.08 |
| **37** | **0.9800** | **0.9093** | **0.8339** | **0.8825** | **0.5311** | **0.7985** | **0.27** |
| 38 | 0.0085 | 0.0020 | 0.0357 | 0.0183 | 0.0000 | 0.0517 | 0.13 |
| 39 | 0.0002 | 0.0007 | 0.0055 | 0.0170 | 0.0000 | 0.0023 | 0.10 |
| 40 | 0.0038 | 0.0014 | 0.0111 | 0.0012 | 0.0000 | 0.0067 | 0.09 |
| 41 | 0.0007 | 0.0005 | 0.0153 | 0.0007 | 0.0000 | 0.0089 | 0.10 |
| 42 | 0.9659 | 0.9917 | 0.8497 | 0.7930 | 0.9334 | 0.8147 | 0.20 |
| 43 | 0.9999 | 0.9999 | 0.9976 | 0.9959 | 1.0000 | 0.9915 | 0.09 |
| 44 | 0.9999 | 0.9999 | 0.9991 | 0.9942 | 1.0000 | 0.9911 | 0.09 |
| 45 | 0.0056 | 0.0015 | 0.0373 | 0.0030 | 0.0000 | 0.0099 | 0.12 |
| 46 | 0.0048 | 0.0053 | 0.0345 | 0.0857 | 0.0000 | 0.0532 | 0.15 |
| 47 | 0.0337 | 0.0293 | 0.0442 | 0.0055 | 0.0000 | 0.0290 | 0.12 |
| 48 | 0.0041 | 0.0200 | 0.0295 | 0.0673 | 0.0000 | 0.0145 | 0.13 |
| 49 | 0.0030 | 0.0120 | 0.0054 | 0.0124 | 0.0000 | 0.0080 | 0.09 |

Here, var\* was calculated using $-1/\log(\text{prediction variance})$ . The bold lines represent “outlier” residues with high CS2BPS prediction variance (whose prediction variance was $> 1.5 \times \text{IQR}$ (the interquartile range)).

## Conflict of interest statement

None declared.

## REFERENCES

1. Guerrier-Takada, C., Gardiner, K., Marsh, T., Pace, N., and Altman, S. (1983) The RNA moiety of ribonuclease P is the catalytic subunit of the enzyme. Cell, 35(3), 849–857.

2. Sharp, P. A. (2009) The centrality of RNA. Cell, 136(4), 577–580.

3. Ponting, Chris P an Oliver, P. L. and Reik, W. (1999) Evolution and functions of long noncoding RNAs. Cell, 136(4), 629–641.

4. Dethoff, E. A., Chugh, J., Mustoe, A. M., and Al-Hashimi, H. M. (2012) Functional complexity and regulation through RNA dynamics. Nature, 482(7385), 322–330.

5. Gesteland, R. F., Cech, T. R., and Atkins, J. F. (1999) The RNA World, Cold Spring Harbor Laboratory Press.

6. Reining, A., Nozinovic, S., Schlepckow, K., Buhr, F., Fürtig, B., and Schwalbe, H. (2013) Three-state mechanism couples ligand and temperature sensing in riboswitches. Nature, 499(7458), 355–359.

7. Haller, A., Rieder, U., Aigner, M., Blanchard, S. C., and Micura, R. (2011) Conformational capture of the SAM-II riboswitch. Nature chemical biology, 7(6), 393–400.

8. In vitro and in vivo studies of the RNA conformational switch in Alfalfa mosaic virus.

9. Cruz, J. A. and Westhof, E. (2009) The dynamic landscapes of RNA architecture. Cell, 136(4), 604–609.

10. Sim, A. Y.-L. and Levitt, M. (2011) Clustering to identify RNA conformations constrained by secondary structure. Proceedings of the National Academy of Sciences, 108(9), 3590–3595.

11. Mathews, D. H., Sabina, J., Zuker, M., and Turner, D. H. (1999) Expanded sequence dependence of thermodynamic parameters improves prediction of RNA secondary structure. Journal of molecular biology, 288(5), 911–940.

12. Mathews, D. H., Disney, M. D., Childs, J. L., Schroeder, S. J., Zuker, M., and Turner, D. H. (2004) Incorporating chemical modification constraints into a dynamic programming algorithm for prediction of RNA secondary structure. Proceedings of the National Academy of Sciences, 101(19), 7287–7292.

13. Mathews, D. H. and H, T. D. (2006) Prediction of RNA secondary structure by free energy minimization. Current opinion in structural biology, 16(3), 270–278.

14. Farés, C., Amata, I., and Carlomagno, T. (2007) ^13^C-detection in RNA bases: revealing structure-chemical shift relationships. Journal of the American Chemical Society, 129(51), 15814–15823.

15. Ohlenschläger, O., Haumann, S., Ramachandran, R., and Görlach, M. (2008) Conformational signatures of 13C chemical shifts in RNA ribose. Journal of biomolecular NMR, 42(2), 139–142.

16. Blad, H., Reiter, N. J., Abildgaard, F., Markley, J. L., and Butcher, S. E. (2005) Dynamics and metal ion binding in the U6 RNA intramolecular stem-loop as analyzed by NMR. Journal of Molecular Biology, 353(3), 540–555.

17. Zhao, B. and Zhang, Q. (2015) Characterizing excited conformational states of RNA by NMR spectroscopy. Current Opinion in Structural Biology, 30, 134–146.

18. Frank, A. T., Law, S. M., and Brooks III, C. L. (2014) A simple and fast approach for predicting ^1^H and ^13^C chemical shifts: toward chemical shift-guided simulations of RNA. The Journal of Physical Chemistry B, 118(42), 12168–12175.

19. Aeschbacher, T., Schubert, M., and Allain, F. H.-T. (2012) A procedure to validate and correct the 13 C chemical shift calibration of RNA datasets. Journal of biomolecular NMR, 52(2), 179–190.

20. Martin, A. D., Quinn, K. M., and Park, J. H. (2011) MCMCpack: Markov chain monte carlo in R.

21. Lu, X. and Olson, W. K. (2008) 3DNA: a versatile, integrated software system for the analysis, rebuilding and visualization of three-dimensional nucleic-acid structures. Nature protocols, 3(7), 1213–1227.

22. Lu, X.-J., Bussemaker, H. J., and Olson, W. K. (2015) DSSR: an integrated software tool for dissecting the spatial structure of RNA. Nucleic Acids Research, 43(21), e142.

23. Burren, S. v. and Groothuis-Oudshoorn, K. (2010) mice: Multivariate imputation by chained equation in R. Journal of statistical software, pp. 1–68.

24. Schneider, G. and Wrede, P. (1998) Artificial neural networks for computer-based molecular design. Progress in biophysics and molecular biology, 70(3), 175–222.

25. Rognes, T., Flouri, T., Nichols, B., Quince, C., and Mahé, F. (2016) VSEARCH: a versatile open source tool for metagenomics. PeerJ, 4, e2584.

26. Merino, E. J., Wilkinson, K. A., Coughlan, J. L., and Weeks, K. M. (2005) RNA structure analysis at single nucleotide resolution by selective 2’-hydroxyl acylation and primer extension (SHAPE). Journal of the American Chemical Society, 127(12), 4223–4231.

27. Wilkinson, K. A., Merino, E. J., and Weeks, K. M. (2006) Selective 2’-hydroxyl acylation analyzed by primer extension (SHAPE): quantitative RNA structure analysis at single nucleotide resolution. Nature protocols, 1(3), 1610.

28. Deigan, K. E., Li, T. W., Matthews, D. H., and Weeks, K. M. (2008) Accurate SHAPE-directed RNA structure determination. Proceedings of the National Academy of Sciences, 106(1), 97–102.

29. Low, J. T. and Weeks, K. M. (2010) SHAPE-directed RNA secondary structure prediction. Methods, 52(2), 150–158.

30. Bellaousov, S., Reuter, J. S., Seetin, M. G., and Mathews, D. H. (2013) RNAstructure: web servers for RNA secondary structure prediction and analysis. Nucleic Acids Research, 41, W471–W474.

31. Mathews, D. H., Disney, M. D., Childs, J. L., Schroeder, S. J., Zuker, M., and Turner, D. (2004) Incorporating chemical modification constraints into a dynamic programming algorithm for prediction of RNA secondary structure. Proceedings of the National Academy of Sciences, 101(19), 7287–7292.

32. Lu, Z. J., Gloor, J. W., and Mathews, D. H. (2009) Improved RNA secondary structure prediction by maximizing expected pair accuracy. Rna, 15, 1805–1813.

33. (2010) ProbKnot: Fast prediction of RNA secondary structure including pseudoknots. Rna, 16, 1870–1880.

34. Hajdin, C. E., Bellaousov, S., Huggins, W., Leonard, C. W., Mathews, D. H., and Weeks, K. M. (2013) Accurate SHAPE-directed RNA secondary structure modeling, including pseudoknots. Proceedings of the National Academy of Sciences, 110(14), 5498–5503.

35. Zou, Q., Xie, S., Lin, Z., Wu, M., and Ju, Y. (2016) Finding the best classification threshold in imbalanced classification. Big Data Research, 5, 2–8.

36. Zhao, B., Guffy, S. L., Williams, B., and Zhang, Q. (2017) An excited state underlies gene regulation of a transcriptional riboswitch. Nature chemical biology, 13(9), 968–974.

37. Marcheschi, R. J., Staple, D. W., and Butcher, S. E. Programmed ribosomal frameshifting in SIV is induced by a highly structured RNA step-loop. Journal of molecular biology, 373(3).

38. Donghi, D., Pechlaner, M., Finazzo, C., Knobloch, B., and Sigel, R. K.-O. (2013) The structural stabilization of the κ three-way junction by Mg(II) represents the first step in the folding of a group II intron. Nucleic Acids Research, 41(4), 2489–2504.

39. Chen, Y., Zubovic, L., Yang, F., Godin, K., Pavelitz, T., Castellanos, J., Macchi, P., and Varani, G. (2016) Rbfox proteins regulate microRNA biogenesis by sequence-specific binding to their precursors and target downstream Dicer. Nucleic acids research, 44(9), 4381–4395.

40. Wu, M. and Tinoco, I. (1998) RNA folding causes secondary structure rearrangement. Proceedings of the National Academy of Sciences, 95(20), 11555–11560.

41. Zhao, B., Hanson, A. L., and Zhang, Q. (2014) Characterizing Slow Chemical Exchange in Nucleic Acids by Carbon CEST and Low Spin-Lock Field R_1ρ_ NMR Spectroscopy. Journal of the American Chemical Society, 136(1), 20–23.

42. Dethoff, E. A., Petzold, K., Chugh, J., Casiano-Negroni, A., and Al-Hashimi, H. M. (2012) Visualizing transient low-populated structures of RNA. Nature, 491(7426), 724–728.

